# *ARID2* deficiency promotes tumor progression and is associated with higher sensitivity to PARP inhibition in lung cancer

**DOI:** 10.1101/2020.01.10.898726

**Authors:** Thaidy Moreno, Laura González-Silva, Beatriz Monterde, Isabel Betancor-Fernández, Carlos Revilla, Antonio Agraz-Doblas, Javier Freire, Pablo Isidro, Laura Quevedo, Santiago Montes-Moreno, Laura Cereceda, Aurora Astudillo, Berta Casar, Piero Crespo, Cristina Morales Torres, Paola Scaffidi, Javier Gomez-Roman, Eduardo Salido, Ignacio Varela

## Abstract

The survival rate in lung cancer remains stubbornly low and there is an urgent need for the identification of new therapeutic targets. Last decade’s research has evidenced a clear role of chromatin structure in cancer development and several members of the SWI/SNF chromatin remodeling complexes have been described altered in different tumor types. Nevertheless, the precise mechanisms of their impact on cancer progression, as well as the application of this knowledge to cancer patient management are largely unknown.

In this study, we have performed targeted sequencing of a cohort of lung cancer patients on genes involved in chromatin structure, as well as functional experiments to identify the molecular mechanisms linking alterations of chromatin remodeling genes and tumor development.

We have identified ARID2 production loss in 20% of lung cancer patients. Additionally, we have shown that ARID2-deficiency provokes profound chromatin structural changes, alters the transcriptional programme and impairs DNA repair which bolster the proliferative and metastatic potential of the cells both *in vitro* and *in vivo*. Moreover, we have demonstrated that ARID2 deficiency significantly affects the sensitivity of the cells to PARP inhibition.

All these results support that *ARID2* is a *bona-fide* tumor suppressor gene in lung cancer that might be exploited therapeutically.

## INTRODUCTION

Lung cancer is the major cause of cancer-related deaths worldwide. The average 5-year survival rate is below 20% irrespective of the subtype, a figure that has only marginally improved in last decades ^1^. Consequently, any new knowledge about the molecular mechanisms that drive this disease could have a great impact on the treatment of patients. In recent years, large genomic projects have facilitated the identification of major players in this tumor type. Thus, small cell lung cancer (SCLC) which constitutes around 15% of all cases, is mainly driven by mutations in *TP53* and *RB1*, but the role of other genes like *PTEN, SLIT2* or *CREBBP* has been also described ^2^. Among non-small cell lung cancer (NSCLC), more than half of the cases are adenocarcinomas, where *KRAS, EGFR, ALK, ROS1* and *BRAF* are the main recurrently altered genes and used as markers for sensitivity to specific anti-tumor therapies ^3^, while squamous cell carcinomas (SCC) are genetically more heterogeneous and poor in actionable mutations so far.

Lately, several members of the SWI/SNF family of chromatin remodeling complexes have been identified recurrently altered in different tumor types adding to the accumulated compelling evidence on the role of chromatin structure in cancer development. It is estimated that approximately 20% of all tumors contain alterations in these complexes, a frequency that is only exceeded by mutations in *TP53* ^4^. In the case of non-small cell lung cancer, the expression of any of the two mutually exclusive catalytic ATPase subunits (SMARCA2 or SMARCA4) is lost in 30% of the cases where it is associated with worse prognosis ^5^. Additionally, *ARID1A*, which encodes for one of the auxiliary subunits of the complex, frequently appears mutated in lung adenocarcinoma ^6^.

## MATERIALS AND METHODS

Detailed protocols can be found in Supplementary Methods.

### Next-generation sequencing

Cancer patient primary samples and, when available, matched corresponding normal samples, were obtained from different tumor Biobanks. In all the cases, we counted with the prior approval of the corresponding ethics committee for each institution. A detailed list of the origin and characteristics of each sample can be found in Supplementary Table 1. DNA was extracted from fresh frozen tissue or cell lines using the Agencourt DNAdvance Beckman Coulter kit (#A48705, Beckman Coulter, Brea, CA, USA), fragmented and submitted to a series of enzymatic steps including End-repair and Adenylation, PE adaptor ligation and PCR indexing amplification. Target capture were performed on pools of 96 libraries using a Sure Select® user-defined probe kit (Agilent Technologies, Palo Alto, CA, USA). Massively parallel sequencing was carried out in a High-Seq® machine (Illumina, USA) with a 100bp paired end (PE) protocol. A single lane was performed for each 96-library pool.

ATAC-Seq libraries were generated according to previously reported protocols with minor modifications. Mainly, cell nuclei were extracted using cold lysis buffer (see supplementary methods) and submitted to tagmentation (Nextera DNA Library Preparation Kit, Illumina, USA). After purification, common adapters included in previous step were used to complete Illumina sequencing full adapters by PCR with Phusion High Fidelity DNA polymerase (Thermo Fisher Scientific, UK). Resulting libraries were submitted for sequencing in a High-Seq machine (Illumina, USA) with a 75 bp paired-end protocol. On average more than 100 million reads were generated in triplicates for each condition.

Total RNA was isolated and purified using Extract Me Total RNA Kit (Blirt, USA). Reverse transcription was performed using the Takara PrimeScript cDNA Synthesis kit (Takara Bio Europe, France). For cDNA library preparations, poly-A mRNA was enriched, fragmented, and submitted to cDNA generation using PrimeScript Enzyme for first strand and RNAse HI, DNA polymerase I and T4 DNA Polymerase (Thermo Fisher Scientific, UK) for the second strand. Afterwards, genomic libraries were generated as above. A minimum of 70 million 75 paired-end reads was generated of 4 replicates of each sample group. Finally, individual mRNA expression was measured by qRT-PCR using Luminaris Color HiGreen qPCR Master Mix (Thermo Fisher Scientific, UK) with StepOnePlusTM real-time PCR system (Applied Biosystems, Foster City, CA). β-actin was used as housekeeping gene and the ΔΔCt method was used for quantification and comparison.

### Sequencing Data Analysis

DNA Raw sequence data was mapped to the human genome (hg19) using BWA 0.7.3 ^7^. Additionally, Samtools 0.1.18 ^8^, Picard 1.61 (http://broadinstitute.github.io/picard/) and GATK 2.2.8 ^9^ was used for format transformation, sorting and indexing of the bam files, as well as to clean the alignment, mark PCR duplicates and perform local realignment around the indels. An in house written software call RAMSES ^10^ and PINDEL 0.2.4d ^11^ were used for substitutions and small insertion and deletion identification respectively. All *ARID2* mutations were validated by PCR amplification coupled with ultrasequencing at a minimum of 10,000 x coverage. Additionally, a similar orthogonal validation of more than 180 mutations randomly picked showed a near 80% of specificity in the mutation calling. Functional consequence of the mutations was annotated using ensembl database v.73 through the Perl API. OncodriveFML software was run to detect genes with evidence of selective pressure from the analysis ^12^.

ATAC-Seq reads were aligned against the human genome (hg19) using BWA 0.7.3 ^7^. Accessible regions were identified using MACS 2.1.2 ^13^. A combined list of all the regions identified in all the samples was used to perform significant region accessibility in ARID2-deficient cells versus control using DESeq2 ^14^. Region annotation was performed using ChIPSeeker software ^15^. BEDTools^16^ was used to estimate the overlapping of the identified regions with ENCODE publicly available data and the results were plotted using deeptools v3.3.1 ^17^. Motif enrichment analysis were performed using HOMER^18^. Finally, alignments were visualized using IGV genome browser ^19^.

Paired-end reads from RNA-Seq were aligned using Tophat ^20^ to the human genome (hg19). Predicted transcripts from Ensembl database were analyzed and transcripts that would lack a CDS start or stop site were filtered out. Differentially expressed genes (DEG) were identified using HTSeq + DESeq ^21,22^.

### Cell Culture and *in vitro* assays

Both A549 and H460 lung cancer cell lines were obtained from The Francis Crick Institute common repository, authenticated by STR profiling, and tested for mycoplasma. For stable cell line generation, tetracycline-inducible pTRIPZ constructs V2THS_74399 (v2), V3THS_347660 (v3) were used for ARID2 knockdown (Dharmacon/GE Healthcare, Lafayette, CO, USA). The empty vector (RHS4750) was used as control. Virus production were performed by transfecting HERK293-T/17clone cells with the pTRIPZ constructs, psPAX2 and pMD2.G plasmids (Addgene) using Fugene HD (Promega Madison, WI, USA). Infected cells were selected with 1 μg/ml puromycin and isolated by FACS using a FACS-Aria II cell sorter (Becton Dickinson, BD, Franklin Lakes, USA) based on TurboRFP expression after the induction with 1 µg/ml of Doxycycline

Growth curve analysis was performed over a period of fourteen days by cell counting with a hemocytometer or by PrestoBlue® assay (Thermo Fisher Scientific, UK). Cell proliferation was also analyzed using the carboxyfluorescein diacetate succinimidyl ester labeling method with the CellTrace™ CFSE Cell Proliferation Kit (Invitrogen, CA, USA) in cells synchronized by gradual serum deprivation following the protocol described by Lauand and collaborators ^23^. The cell proliferation index was analyzed using MODFIT software (Verity, USA). Proliferation index was the sum of the cells in all generations divided by the calculated number of original parent cells.

*In vitro* cell migration assays were performed by using 8-μm pore size transwell chambers (Corning™ Transwell™ Multiple Well Plate with Permeable Polycarbonate Membrane Inserts, ref 3422) in 24-well plates using 10% FBS in the lower chamber served as the chemo-attractant. Filters were fixed in 4% PFA followed by crystal violet staining for microscope visualization. For the invasion assays, the cells were plated on a growth factor-reduced Matrigel (BD Biosciences) pre-coated 8 μm pore transwell chamber. Invasive cells were quantified by fixing chambers in 4% paraformaldehyde for 10 min and staining with crystal violet.

Inhibition assays were performed to determine the half maximal inhibitory concentration (IC50) values for different antitumoral drugs. Cells were cultured for 24 hours in different drug concentrations. Viability was determined by PrestoBlue® reagent (Thermo Fisher Scientific, UK) and measured using a Multiskan FC Microplate Photometer (Thermo Fisher Scientific, Waltham, MA) with wavelengths set at 540 and 620 nm. IC50 value for each drug were determined with Prism software (GraphPad, USA) to fit curves to the dose response data.

### *In vivo* tumorogenesis assays

Animal studies were conducted in compliance with guidelines for the care and use of laboratory animals and were approved by the Ethics and Animal Care Committee of the Universidad de Cantabria. For proliferation assays, five million cells were subcutaneously injected into the flanks of the 6-8-week-old female nude mice (Athymic Nude-Foxn1nu, Envigo, UK). For metastasis assays, 2.5 million of cells were tail injected into 6-8-week-old female nude mice. The animals were treated with 2 mg/mL of Doxycycline in the drinking water.

### Western blot analysis

Cells were washed twice in PBS and lysed in RIPA buffer (50 mM Tris-HCl, pH 8.0, 150 mM NaCl, 1 % NP-40, 1 mM Sodium Orthovanadate, 1 mM NaF) containing Halt protease inhibitors Cocktail (Thermo Scientific, ref 87786). Lysates were sonicated using the Bioruptor® (Dia-genode) and cleared by centrifugation at 16,000g for 20 min at 4 °C. Total protein lysates were separated by SDS-PAGE in 8% polyacrylamide gels and transferred to nitrocellulose membranes. Subsequently, membranes were washed with TBS-T (50 mM TRIS + 150 mM Sodium chloride + 0,1% Tween 20, pH 7,4) and blocked using 5% non-fat milk solution as blocking agent in TBS (50 mM TRIS + 150 mM Sodium chloride) for 1 h at RT. Membranes were then incubated with primary antibodies anti-ARID2 (E-3, sc-166117, Santa Cruz) and anti-Actin (I-19, sc-1616, Santa Cruz), diluted 1:200 and 1: 1,000 in TBS-T/5% (w/v) BSA at 4°C overnight, respectively. Donkey anti-mouse or donkey anti-goat secondary antibodies (LI-COR Biotechnology, Lincoln, USA) conjugated to IRDye 800CW (926-32212) or IRDye 680RD (926-68074) respectively were used as secondary antibodies and visualized using Odyssey Clx imager (LI-COR Biotechnology, Lincoln, USA).

### Immunohistochemistry analysis

For ARID2 detection on paraffin sections, antigen retrieval was performed for 32 minutes at 97 °C in citrate buffer pH 6, incubated with 1:300-1:500 anti-ARID2 antibody (abcam ab113283) and developed with HRP-polymer secondary antibodies (Optiview, Roche). ARID2 expression was evaluated by two pathologists on coded tissue sections, without information about the *ARID2* mutation status. Only surgical pathology cases with enough material, both tumor and non-neoplastic surrounding tissue, were considered for ARID2 immunohistochemistry. After initial independent evaluation by the two pathologists, scoring ARID2 expression relative to normal adjacent tissue, a consensus score was reached viewing the slides at a multiheaded scope.

Immunofluorescence was performed in stable cells fixed with 4% paraformaldehyde in PBS for 15 min at room temperature. The cells were permeabilized with 0.5% Triton X-100 in PBS and blocked with 3% BSA in PBT (PBS containing 0.05% Triton X-100). Finally there were subjected to immunofluorescence staining with ARID2 antibody (E-3, sc-166117 Santa Cruz, CA, USA or A302-230A, Bethyl Laboratories, USA), anti-phospho-Histone H2A.X Ser139 (γH2AX, clone JBW301, Merck Millipore, USA) or anti-53BP1 antibody (H-300, sc-22760, Santa Cruz, USA) Cover slides were incubated with Alexa labeled secondary antibodies and mounted in VECTASHIELD Antifade Mounting Medium with DAPI (Vector Labs, USA). The cells were finally examined by fluorescence microscopy (Olympus America Inc, Center Valley, PA or Nikon Eclipse 80i upright fluorescence microscope Tokyo, Japan). Colocalization of ARID2 with 53BP1 or γH2AX was performed measuring the variation in intensity across the lines drawn using the linescan tool from MetaMorph® (Molecular Devices, CA, USA). Automatic foci quantification was performed in randomly acquired images, modified the speckle counting pipeline from CellProfiler open-source software ^24^.

### Data availability

DNA-Seq, ATAC-Seq and RNA-Seq data: European Nucleotide Archive PRJEB26936 https://www.ebi.ac.uk/ena/data/search?query=PRJEB26936

## RESULTS

### Loss of ARID2 protein expression in 20% of lung cancer patients is partially explained by the presence of driver *ARID2* mutations

In order to understand better the role of chromatin remodeling complexes in lung cancer development, we performed a genetic screening on the coding sequences of known cancer genes as well as members of the main chromatin remodeling complexes (Suppl. Table 2). We applied targeted next-generation sequencing technologies in a collection of 81 lung cancer cases (40 lung adenocarcinomas, 12 squamous cell carcinomas and 29 small cell carcinomas) (Suppl. Table 1). Interestingly, we found mutations in *ARID2* in 5 of the patients (Figure 1a and Suppl. Table 3). Additionally, when we run OncodriveFML ^25^ analysis to identify lung cancer driver genes in our data, *ARID2* showed significant positive selection after multi-test correction, just below *TP53* (Figure 1b and Suppl. Table 4). To validate these results, we performed targeted sequencing in a second cohort of 144 lung adenocarcinoma cases finding mutations in 12 patients, for a total frequency of *ARID2* mutations of 7.5 % (17/226) (Suppl. Table 3). In the case of lung adenocarcinoma, we found *ARID2* non-synonymous mutations in 7% of the samples (13/185) which is higher than the frequency reported in COSMIC database for this tumor type (3.7 %, 83/2241) ^26^ and ranks *ARID2* among the ten genes most commonly mutated in lung cancer. In concordance with a potential role of *ARID2* as tumor suppressor, many of the identified mutations, clustered at the beginning of the protein sequence, are predicted to generate a premature truncation of the protein (Figure 1c and Suppl. Table 3). Subsequently, to check if the loss of ARID2 function is a common feature in lung cancer, we performed immunohistochemistry analyses in 139 of the studied samples finding loss or low/heterogeneous ARID2 production in approximately 20% of the cases (28/139) (Figure 1d and Suppl. Figure 1). Additionally, loss of *ARID2* expression is significantly more frequent in *ARID2*-mutated patients (6/10 Fisher exact test p=0.0098). Interestingly, this is also true for some samples with missense mutations, which suggests that these mutations might interfere with the correct folding or processing of the protein. Indeed, many of the mutations found are predicted to produce deleterious effects in the protein according to SHIFT or Polyphen algorithms (Suppl. Table 3). The presence of normal tissue contamination in the tumor samples as well as the sequencing strategy followed for normal tissue samples, prevented an estimation of the cellularity or the zygosity of the mutations to check if the wildtype *ARID2* allele is lost during tumor progression. The fact that some non-mutated samples show also loss of ARID2 production suggests the existence of other, maybe epigenetic, mechanisms that could interfere with *ARID2* expression.

**Figure 1.**
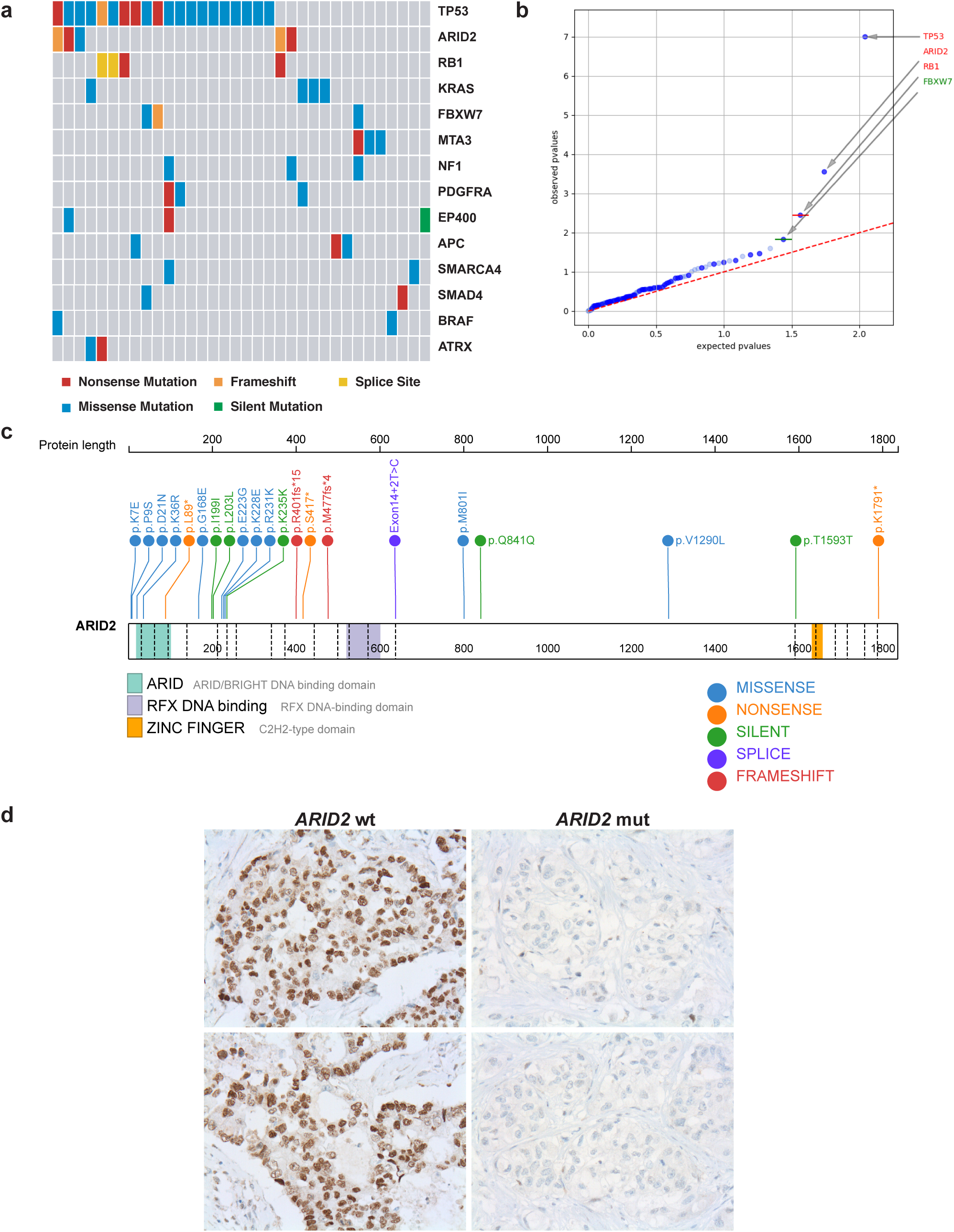
Frequent ARID2 mutations associated with loss of protein synthesis and worse prognosis in lung cancer patients. (a) Box representation of the mutated patients for the most significantly mutated genes according to OncodriveFML in the lung cancer cohort generated with Maftools ^48^. Each box in the central matrix represents an independent patient. Colored boxes represent mutated patients for the corresponding gene in a color code indicating the type of mutation. (b) Representation of the significance analysis of the functional impact of the mutations found in each gene performed by OncodriveFML. Genes in read showed a q-value < 0.1 after multi-test correction. (c) Visual representation of the location of all identified *ARID2* mutations in our discovery and validation lung cancer cohort in relation to the functional protein domains. (d) Representative images of ARID2 immunohistochemistry experiments in two *ARID2*-mutated (right) and two *ARID2*-wildtype (left) lung adenocarcinomas.

### ARID2-deficiency increases proliferative and metastatic potential *in vitro* and *in vivo*

In order to check if alterations in *ARID2* could promote lung cancer development, we performed knock-down experiments *in vitro* using shRNAs. As it can be observed in Figure 2 a-b, *ARID2* mRNA and protein production was efficiently reduced by two different shRNAs. This reduction was accompanied by an increase on the proliferation of A549 lung adenocarcinoma cell line. Additionally, we observed increased invasion and migration capacities of ARID2-deficient A549 and H460 NSCLC cell lines compared to those cells transduced with the empty vector (Figure 2 c-e and Suppl. Figure 2). Moreover, when these cells were injected on immunocompromised mice, they showed a greater capacity to produce tumors *in vivo* (Figure 2f and Suppl. Figure 3).

**Figure 2.**
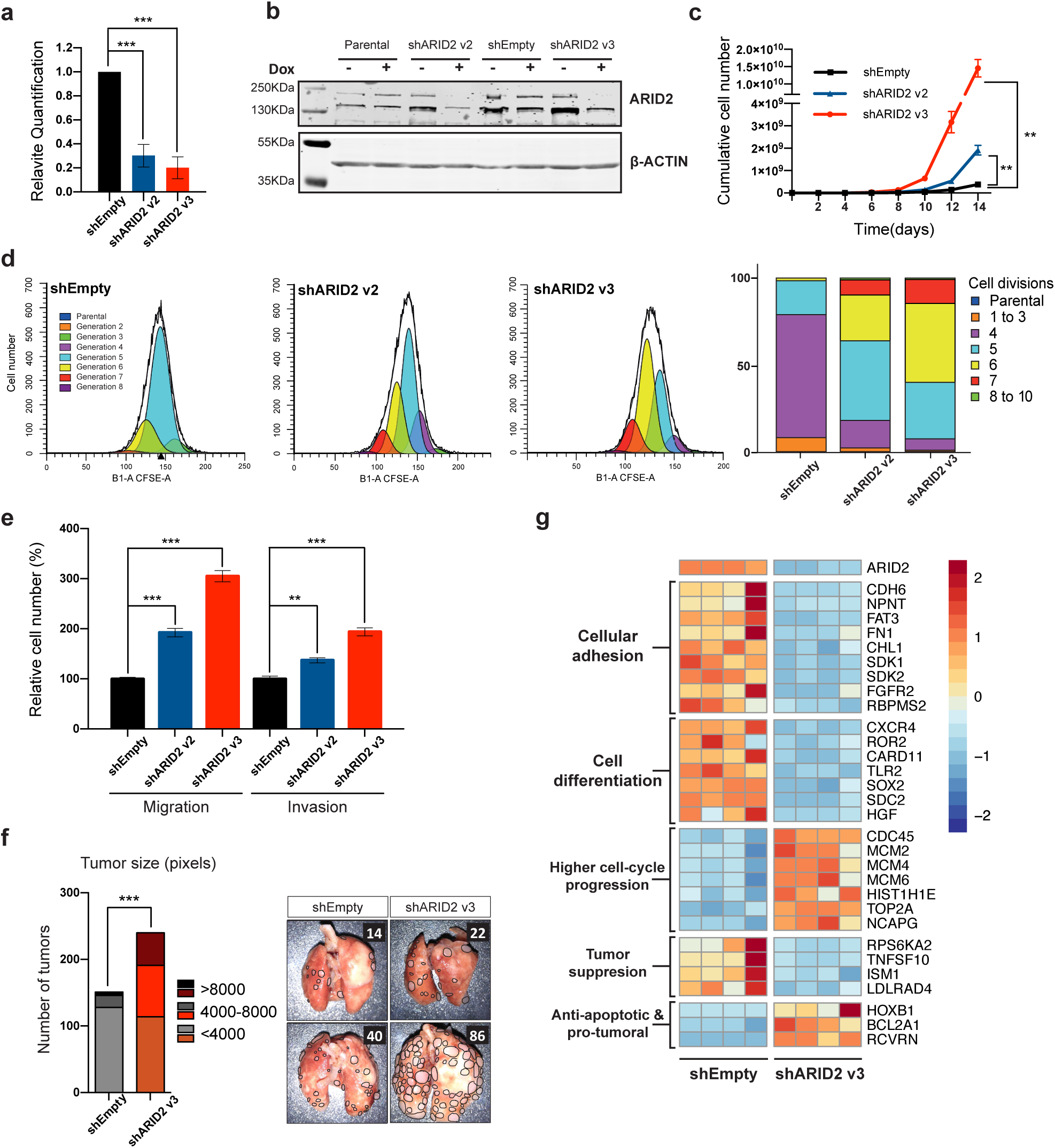
ARID2 deficiency is associated with an increase in oncogenesis *in vitro* and *in vivo*. (a) Bar representation of ARID2 expression level fold changes measured by qRT-PCR in A549 cells transduced with shARID2 v2 and v3 as well as the empty vector which is used as control. (b) Representative image of a western blot analysis measuring ARID2 protein levels in A549 parental cells as well as those cell lines transduced with ARID2 shRNAs and the empty vector. In all the cases, the results are shown with and without induction of the shRNA expression by doxycycline (Dox) treatment. (c) Proliferation curves showing the calculated cell number accumulated by serial cell passaging in empty and ARID2 shRNAs transduced A549 cell lines. (d) Representative experiment of the number of cell divisions suffered by the cells in 48h estimated by CFSE labelling in A549 cells by flow cytometry. Bar quantification on the number of cells that have suffered each number of cell division is represented on the right. (e) Bar representation of quantified cells in destination changer on migration and invasion assays of A549 cells transduced with two different ARID2 shRNAs (blue and red bars). Data shown as mean ± SEM of three independent experiments, relative to control cells A549 Empty vector (black bars). (f) Representative images of lung metastasis generated in intravenously injected mice with A549 cells transduced either with shEmpty, or shARID2 v3 vectors. Individual metastasis are delineated in the image and counted (upper right corner numbers). On the left, a quantification of the number and size of the tumors generated in the two groups is shown. (Fisher exact test *p < 0.05, ** p < 0.01 and *** p <0.001). g) Heatmap representation of a selection of differently expressed genes in ARID2-deficient cells (n=4) and grouped according to their biological function. Expression differences goes from red (overexpression) to blue (downregulation) according to the log2 of the fold change.

Finally, RNA-Seq experiments in the transduced cell lines showed that loss of ARID2 was accompanied with changes in gene expression that support the observed phenotypes in the cells (Figure 2g and Suppl. Table 5). Thus, we observed a downregulation of genes involved in cellular adhesion and cell differentiation such as *NPNT, CDH6, FAT3, FN1, SOX2* or *SDC2* as well as an upregulation of genes associated with a higher cell-cycle progression such as *CDC45, MCM2* or *HIST1H1E*, which could be associated with the increased proliferation, migration and invasion capacities of ARID2-deficient cells. Additionally, we observed downregulation of other tumor suppression genes like *RPS6K2, TNFSF10, ISM1* or *LDLRAD4* together with upregulation of protumoral and anti-apoptotic genes like *HOXB1, BCL2A1* or *RCVRN*. Transcriptional changes in selected genes were further validated by qRT-PCR in independently generated ARID2 knock-down cell lines (Suppl. Figure 4).

Altogether, these results prove that *ARID2* plays a tumor suppressor function in lung cancer.

### ARID2-deficiency is accompanied with profound chromatin changes around enhancers and promoters

We hypothesized that gene expression changes observed in ARID2-deficient cells might by the result of changes in SWI/SNF chromatin remodeling activity. To investigate this, we performed ATAC-Seq experiments in transduced cells. ARID2 loss is accompanied with a significant increase of chromatin accessibility in 687 regions (Suppl. Table 6). These regions were located near of transcription start sites and showed higher enrichment of H3K4me3 histone mark according to ENCODE project data from A549 cell line, revealing the prevalence of promoter regions (Figure 3a). This indicates that ARID2 plays a repressive role in specific genes, keeping a closed chromatin structure around their promoters. On the other hand, ARID2-deficient cells show a significant loss of chromatin accessibility in 990 regions (Suppl. Table 6). These regions were located distal to gene transcription start sites (Figure 3b) and showed enrichment of enhancer specific H3K4me1 and H3k27ac histone marks. Concordant with this, AP-1 family transcription factor binding motif, described as abundantly present in enhancers^27^, is highly enriched on those regions that showed less accessibility on ARID2-deficient cells (Figure 3c). All this support that *ARID2* is essential to keep an open chromatin conformation around selected enhancers.

**Figure 3.**
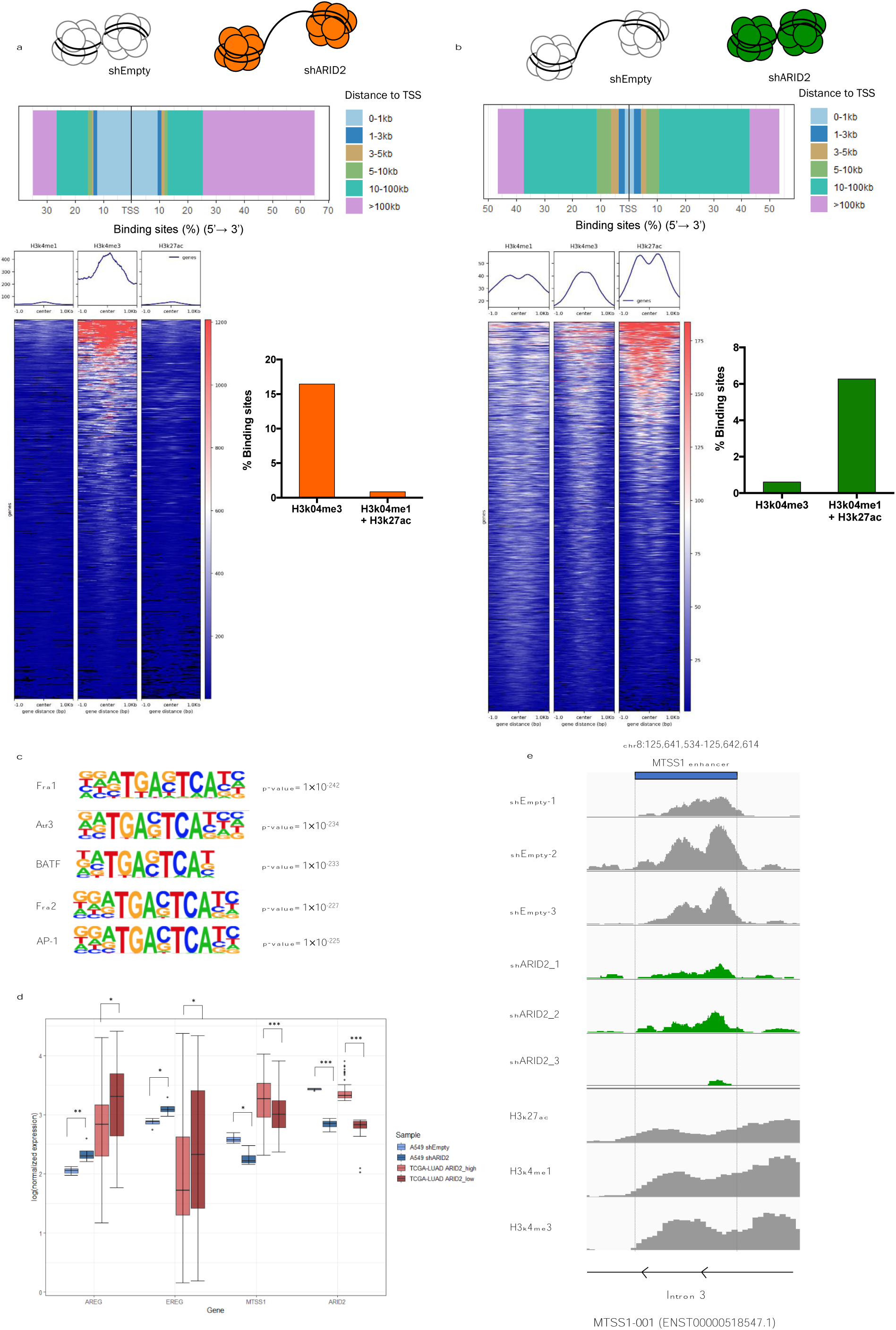
Profound chromatin structural changes affect gene expression after ARID2 loss. a) Analysis of the genomic regions that gain significantly chromatin accessibility after ARID2 loss. In the upper pannel the regions are grouped according to their distance to nearest gene transcription start site (TSS). Below, the intensity of H3K4me1, H3K4me3 and H3k27ac histone marks in each identified region is represented by heatmaps (left). Additionally, the percentage of identified regions that overlap with regions with histone modification marks compatible with active promoters (H3K4me3) or enhancers (H3K4me1 + H3K27ac) are represented in a bar graph (right). b) Analysis of the genomic regions that loose significantly chromatin accessibility after ARID2 loss. In the upper pannel the regions are grouped according to their distance to nearest gene transcription start site (TSS). Below, the intensity of H3K4me1, H3K4me3 and H3k27ac histone marks in each identified region is represented by heatmaps (left). Additionally, the percentage of identified regions that overlap with regions with histone modification marks compatible with active promoters (H3K4me3) or enhancers (H3K4me1 + H3K27ac) are represented in a bar graph (right). c) Enrichment of sequence motifs identified in those regions that gain chromatin accessibility after ARID2-loss. d) Boxplot graph of gene expression differences identified in both our ARID2-deficient cells and in lowly *ARID2* expressing lung adenocarcinoma patients (ARID2_low) versus highly *ARID2* expressing patients (ARID2_high) from TCGA database. e) Visualization, in a described MTSS1 internal enhancer, of read alignments for the different replicates of our ATAC-Seq experiments in ARID2-deficient A549 cells. Additionally, read alignments of ChIP-Seq experiments performed against different histone marks during ENCODE project are also represented.

Altogether, these results indicate that ARID2 plays a dual regulatory role. On one side, ARID2 is essential to keep an open chromatin structure around the enhancers of positively regulated genes. On the other, ARID2 containing complexes regulate negatively specific genes maintaining a closed inactive chromatin structure around their promoters.

### ARID2 is essential to maintain the expression of the metastasis inhibitor *MTSS1*

In order to identify those genes whose expression might be deregulated specifically as the result of ARID2 loss, we compared our RNA-Seq results with differential expression analysis performed on human lung adenocarcinoma patients from TCGA database. Eighteen genes were found upregulated in both ARID2-deficient cells, and in low-ARID2 expressing lung adenocarcinoma patients (Suppl. Table 7). Among them, we found *AREG, ERG* or *NGF* growth factors that might explain the higher proliferation capabilities of ARID2-deficient cells (Figure 3d and Suppl. Table 7). In addition, we found 133 genes downregulated in both datasets indicating a main gene expression activating role of ARID2 in this cellular context.

Interestingly, among those genes whose expression rely on ARID2, we found *MTSS1*, a well described metastasis inhibitor ^28,28,29^. The observed significant reduction of *MTSS1* expression likely explain the higher invasion capabilities of ARID2-deficient cells. Additionally, we found a significant reduction of chromatin accessibility on an intronic region annotated as an internal *MTSS1* enhancer in geneHancer database after ARID2 loss ^30^. This observation is concordant with the hypothesis that *MTSS1* expression is positively regulated by ARID2 by keeping an open chromatin structure at its enhancer (Figure 3e).

### ARID2 co-localizes to DNA-repair foci and its deficiency is associated with DNA-damage accumulation

Gene-set enrichment analyses (GSEA) on the transcriptional alterations observed in ARID2-deficient cells also showed a significant upregulation of genes involved in DNA repair, which reflects an activation of DNA detection and repair mechanisms in these cells (Figure 4a). Consequently, we hypothesized that *ARID2* might play important roles in DNA repair. In order to explore this possibility, we performed immunofluorescence experiments to check the nuclear location of ARID2 in cells subjected to DNA damage. We observed that ARID2 co-localizes with γH2AX and 53BP1 at the DNA repair foci (Figure 4b and Suppl. Figure 5). Moreover, ARID2-deficient cells showed an increase in the number of DNA damage foci in response to DNA damaging agents (Figure 4 c,d). All this demonstrates that ARID2 deficiency interferes with DNA-repair mechanisms unveiling a second tumor promoting mechanism whereby ARID2 deficiency promotes carcinogenesis.

**Figure 4.**
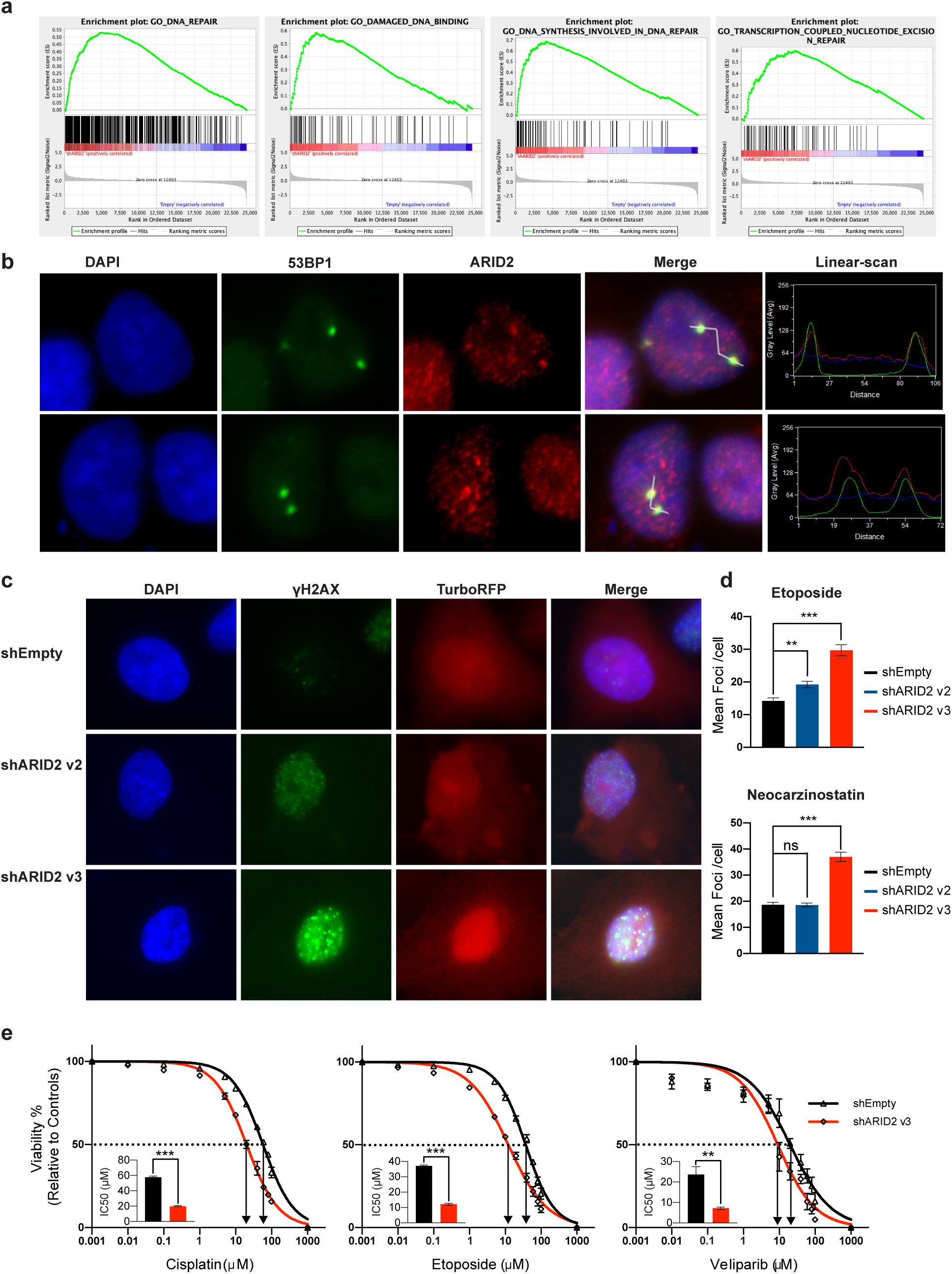
ARID2 deficiency affects DNA repair and affects sensitivity to anti-tumor therapies. (a) Results of the Gene-Set Enrichment Analysis (GSEA) showing enrichment of genes involved in different DNA repair ontologies. (b) Representative images of immunofluorescence experiments demonstrating colocalization of 53BP1 (green) and ARID2 (red) in A549 cells treated with Neocarzinostatin. Colocalization is confirmed on the right through the parallel quantification of red and green signals on a manually selected path through the image using the LineScan tools from Methamorph software (c) Representative images of DNA-repair foci visualized by H2AX immunofluorescence (green) in transduced (red) cells after the treatment of DNA-damaging agents. (d) Bar representation of the foci quantification in each transduced cell line. (e) Representative experiments measuring cell survival to increasing concentrations of cisplatin, etoposide and veliparib on A549 cells transduced with shEmpty (black), or shARID2v3 (red) vectors. Bar grapths represent the calculated IC50 value for each experiment. In all cases, the results are represented as mean +/- SEM of three independent results, (two-tailed t-test *p < 0.05, ** p < 0.01 and *** p <0.001).

### ARID2-deficiency increases cell sensitivity to chemotherapy and veliparib

Considering the observed defects in DNA repair, we studied the possibility of exploiting ARID2 deficiency for cancer treatment. As platinum-based chemotherapy is widely used for the treatment of lung cancer patients ^3^, we studied ARID2-deficient cell lines sensitivity to these therapies. As it can be seen in Figure 4e and Suppl. Figure 6, in concordance with a role of ARID2 in DNA repair, ARID2-deficient cells exhibited a higher sensitivity to DNA damaging agents like cisplatin or etoposide compared to controls.

Additionally, in the last decade, many researchers have described a higher sensitivity of PARP inhibitors in tumors harboring defects in DNA repair mechanisms due to synthetic lethality ^31^. Consequently, we checked if this might apply as well to ARID2-deficient cells. As it can be seen in Figure 4e, ARID2 loss is accompanied with a higher sensibility to veliparib, a well described PARP inhibitor that is under research in several clinical trials in breast, ovarian and, most importantly, lung cancer. Therefore, ARID2 deficiency could potentially be used as marker to direct PARP inhibitor treatment to lung cancer patients.

## DISCUSSION

Although some evidence of the presence of *ARID2* alterations in lung cancer have been reported previously ^32^, the relevance of these alterations for oncogenesis has not been clearly proved. Our results showed an *ARID2* mutation recurrency higher than the one reported in COSMIC database. Additionally, the distribution and predicted impact of the mutations found and our *in-vitro* and *in-vivo* experiments provided compelling evidence of the role of *ARID2* as *bona-fide* tumor suppressor in lung cancer. Supporting this, *ARID2* has been already proposed as cancer driver gene in melanoma and hepatocellular carcinoma ^33,34^.

The precise molecular mechanisms by which alterations in chromatin remodeling complexes promote cancer development are not sufficiently understood. Interactions with well-described cancer genes like *TP53, RB* or *MYC* have been described ^35–37^. In addition to this, they play essential roles in the activation of differentiation and the suppression of proliferative programs of many cellular lineages ^38^. In this study we described a list of near 200 genes that are specifically deregulated after ARID2 loss in both our cellular model and in lung adenocarcinoma patients from the TCGA database. Some of these genes, like *AREG, EREG* or *NGF* growth factors might account for the higher proliferation capabilities of ARID2-deficient cells. In terms of the molecular mechanisms behind this regulation, we show that ARID2-deficiency is associated with profound chromatin structural changes. Our results prove that ARID2 is essential to keep an open chromatin structure in enhancer regions in agreement with an important role of different SWI/SNF members in regulating enhancer activity ^39,40^. One of these ARID2-dependent enhancers regulates MTSS1 expression that, consequently, shows a significant downregulation in ARID2-deficient cells. This suggests that ARID2 might regulate directly *MTSS1* expression, although further work is necessary to finally confirm this. MTSS1 is a well described migration and invasion inhibitor, associated with worse prognosis in several tumor types ^28,29,41^ and its deficiency might well explain the higher migration and invasion capabilities of ARID2-deficient cells.

As a complementary mechanism, we observed an active role of ARID2 in the detection and repair of DNA damage *in vitro* in lung cancer cell lines. In accordance with this view, other members of the SWI/SNF complex have been shown to be involved in different steps of DNA damage repair ^42–44^.

Finally, any advance in the possibility of exploiting therapeutically any vulnerability associated to deficiency in SWI/SNF complex genes is of great interest, as approximately 20% of all human cancers are reported to have alterations in this complex. In this study ARID2-deficient cells showed a higher sensitivity to different DNA-damaging therapies, likely as a result of the ARID2 involvement in DNA repair. Considering that platinum-based chemotherapy is still widely used in lung cancer patients with high variable success ^3^, our results suggest that ARID2 expression might be explored as potential response biomarker for these therapies. Moreover, we show that ARID2 deficiency shows synthetic lethality with PARP inhibition using veliparib, an inhibitor that has shown good results in the treatment of breast cancer ^45^ and is included in several clinical trials on breast, ovarian and, most importantly, lung cancer. Our results suggest that the stratification of lung cancer patients according to ARID2 expression might improve the efficiency of PARP inhibitors in non-small cell lung cancer. Additionally, a very recent study has shown that ARID2-deficient melanoma cells are particularly sensitive to immunotherapy through alterations in mTORC1 and IFNγ pathways ^46^. Interestingly, we observed that some downstream response genes in these pathways such as *GBP2, GBP3* and *SCD5* are significantly downregulated in ARID2-deficient cells (Suppl. Table 5). All these results support the potential use of *ARID2* expression as a new biomarker for personalized treatment in lung cancer patients.

In summary, here we present compelling evidence for the role of *ARID2* as tumor suppressor in lung cancer. Although *ARID2* has been proposed as a driver gene in other tumor types ^34,47^, little has been reported about the molecular mechanisms underlying this involvement. In this work, we propose that its role in lung cancer is exerted in two ways, firstly by fostering a specific pro-oncogenic transcriptomic program as a result of changes in chromatin structure, and secondly by impairing DNA repair. Importantly, our results indicate that ARID2-deficiency could be exploited for lung cancer patient treatment.

## ACKNOWLEDGEMENTS

I.V. is supported by SAF2012-31627 and SAF2016-76758-R grants from the Spanish Ministerio de Economía y Competitividad (MINECO), by a Fundación Ramón Areces grant and by ERC2014-StG637904 grant from the European Research Council. I.V has been awardee of the Programa Ramón y Cajal (MINECO, Spain). T.M has been awardee of the Ayudas para la contratación de investigadores predoctorales (MINECO, Spain). PC laboratory is supported by grant SAF-2015 63638R (MINECO/FEDER, UE); by Centro de Investigación Biomédica en Red de Cáncer (CIBERONC) and by Asociación Española Contra el Cáncer (AECC), grant GCB141423113. BC is supported by a Retos Jóvenes Investigadores grant SAF2015-73364-JIN (AEI/FEDER, UE) and a grant from Fundación Francisco Cobos. P.S. is supported by the Francis Crick Institute, which receives its core funding from Cancer Research UK (FC001152), the UK Medical Research Council (FC001152), and the Wellcome Trust (FC001152). We would like as well to acknowledge the support of the Servicio Santander Supercomputación, the technical staff of IBBTEC and the Servicio de Estabulación and Experimentación Animal (SEEA-UC). We want to thank to Jose Pedro Vaqué and Javier Leon labs (IBBTEC) for providing the different anti-tumor therapies. We want to thank also Francisco Real for their careful review of this manuscript and their advice during the project. Finally, we would like to thank all patients that have agreed to participate in this study and the work of the staff members of the different tumor biobanks that manage patient samples, specially to the Biobanco Valdecilla (Tissue Node, PT13/0010/0024) and the Principado de Asturias BioBank (PT17/001570023) HUCA/IUOPA which is jointly financed by Servicio de Salud del Principado de Asturias, Instituto de Salud Carlos III and Fundación Bancaria Cajastur and integrated in the Spanish National Biobanks Network, for their exceptional work in sample collection and organization.

## AUTHOR CONTRIBUTIONS

T.M., L.G.-S., C.R., L.Q., B.C., C.M.T, P. S., P.C., and I.V. performed the sequencing, molecular, cellular biology and animal experiments and contributed to the analysis of the data. T.M., A.A.-D. and I.V. performed the analysis of next-generation sequencing data. I. B.-F., J. F., P. I., S. M.-M., L. C., A. A., E. S. and J. G.-R. collected and checked the human samples and performed the immunohistochemical analysis. P. C., P. S., E. S. and I. V. contributed in the design and supervision of the experiments and in the elaboration of the manuscript.

## CONFLICT OF INTEREST

The authors disclose no potential conflicts of interest.

## FIGURE LEGENDS

**Supplementary Figure 1.**
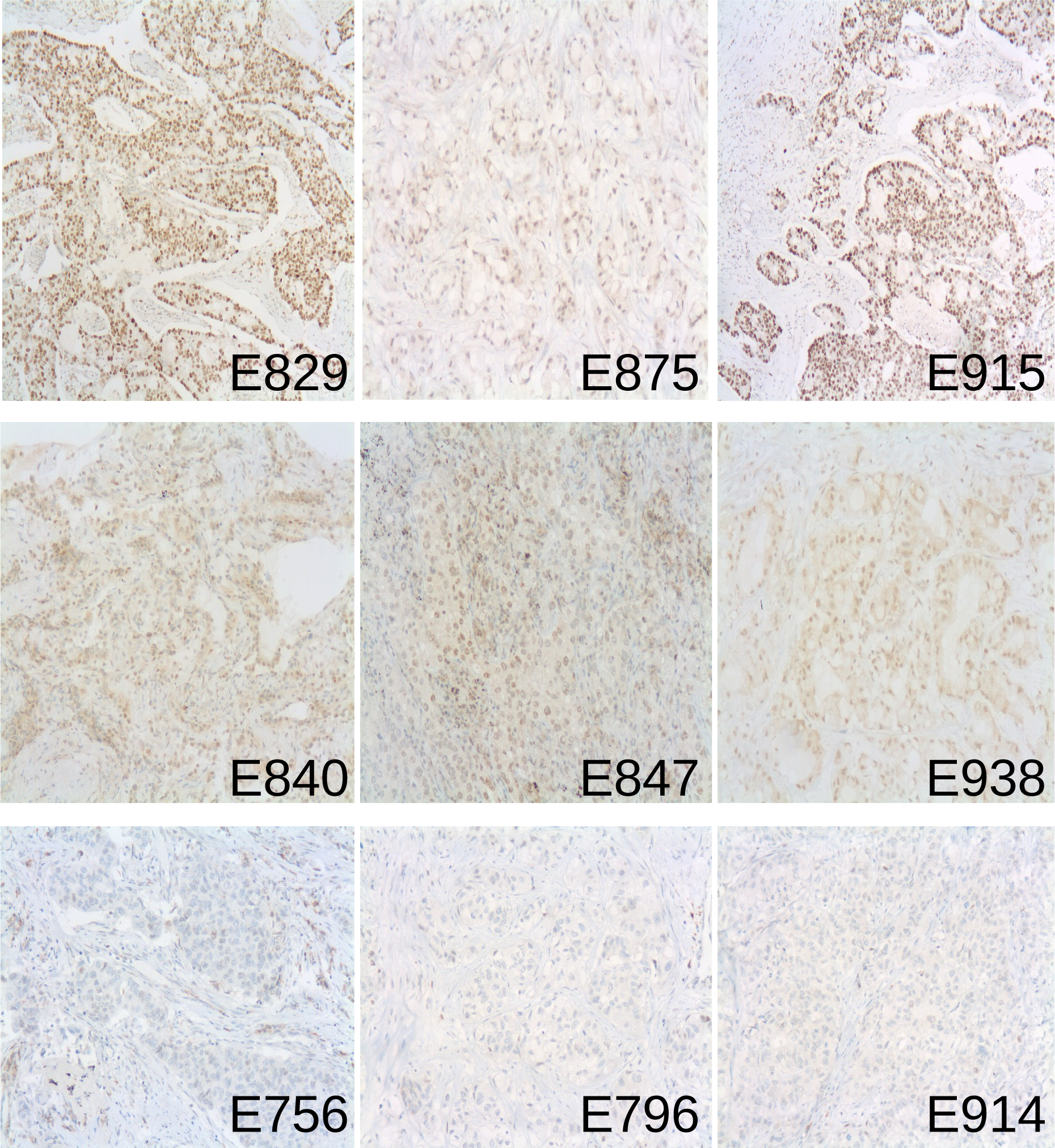
ARID2 mutations affect protein synthesis. ARID2 expression in representative lung adenocarcinomas from the validation cohort. Strong nuclear immunostaining, clearly restricted to the nuclei are shown in the upper panel, while other cases show moderate ARID2 expression (middle panel), not only on the nuclei but also with cytoplasmic background. The bottom row represents adenocarcinomas lacking ARID2 immunostaining.

**Supplementary Figure 2.**
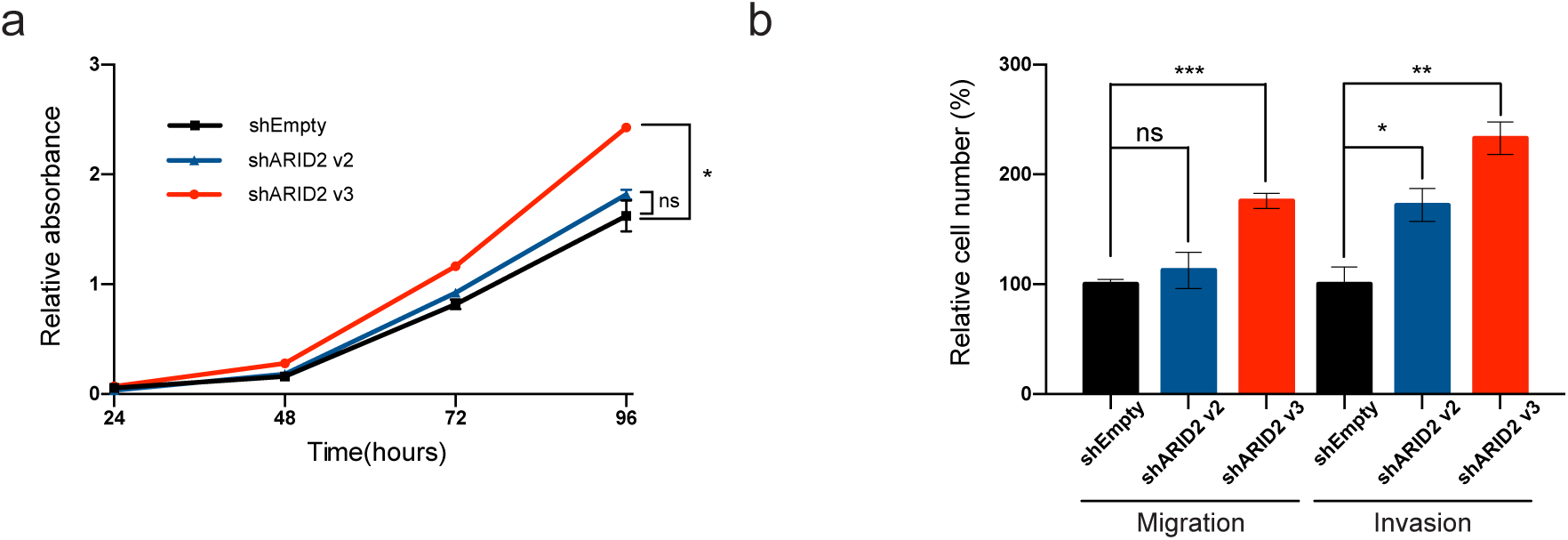
ARID2 deficiency is associated with increase proliferation, migration and invasion *in vitro*. (a) Proliferation curves of A549 transduced cells measured by Preston Blue metabolic labeling. (b) Bar representation of the quantification of the number of cells of the lower chamber in migration and invasion *in vitro* assays in H460 cell lines transduced with empty and ARID2 shRNAs v2 and v3 vectors. (one-tailed t-test *p < 0.05, ** p < 0.01 and *** p <0.001).

**Supplementary Figure 3.**
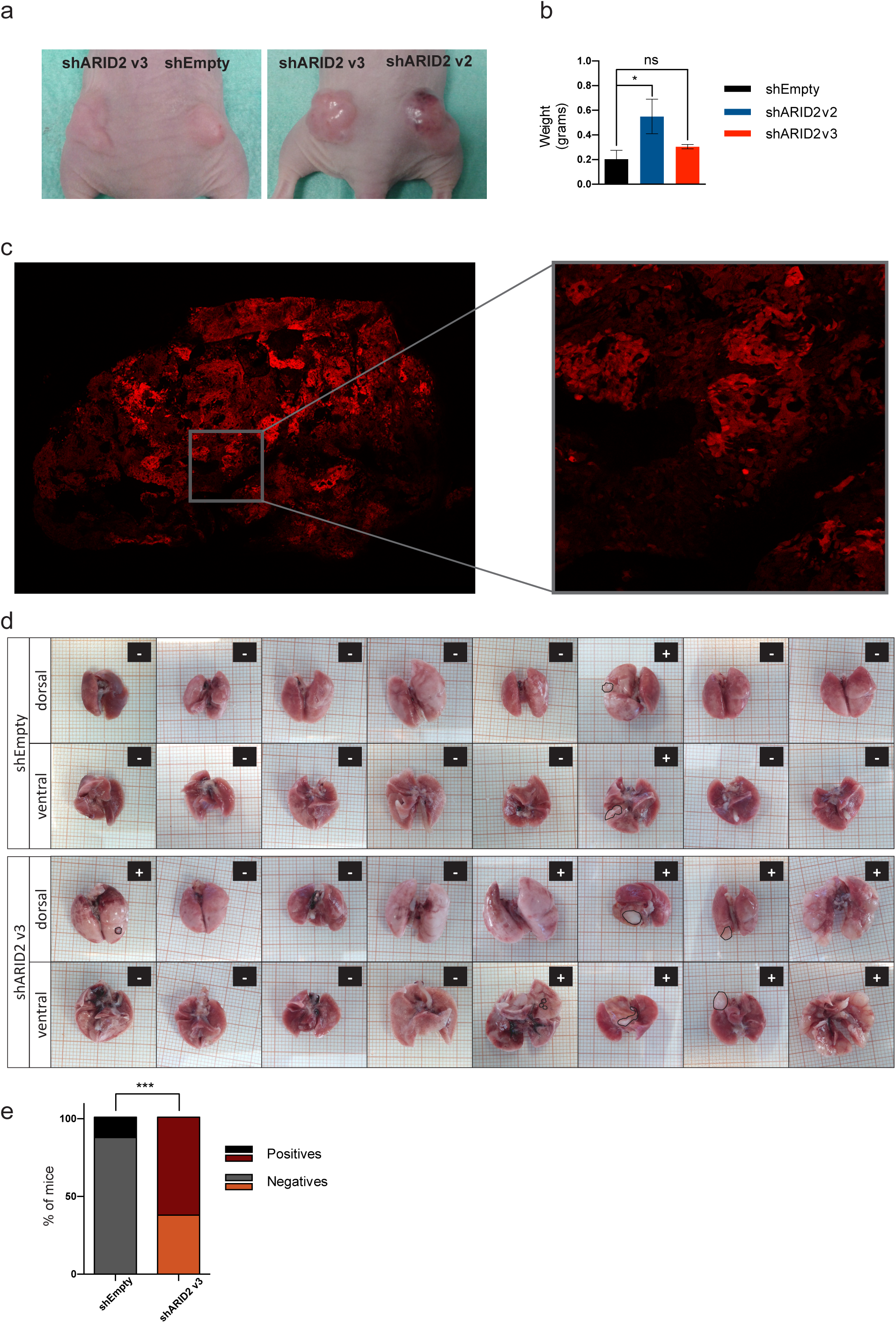
ARID2-deficient cells show and increase in oncogenic potential *in vivo*. (a) Representative images of tumors generated in the flanks of immunocompromised mice injected with A549 cells transduced with empty and ARID2 shRNA vectors. (b) Bar representation of the quantification of the weight of the generated tumors. The results are represented as mean +/- SEM (c) Representative image of a fluorescence analysis of the expression of TurboRFP reporter protein of nude mice xenograft tumors created by A549 stable cells. (d) Images of the tumors generated in the lung of tail-vein injected immunocompromised mice with transduced H460 cell lines. (e) Bar representation of the percentage of mice with tumors. (one-tailed t-test in a, and fisher exact test in e *p < 0.05, ** p < 0.01 and *** p <0.001)

**Supplementary Figure 4.**
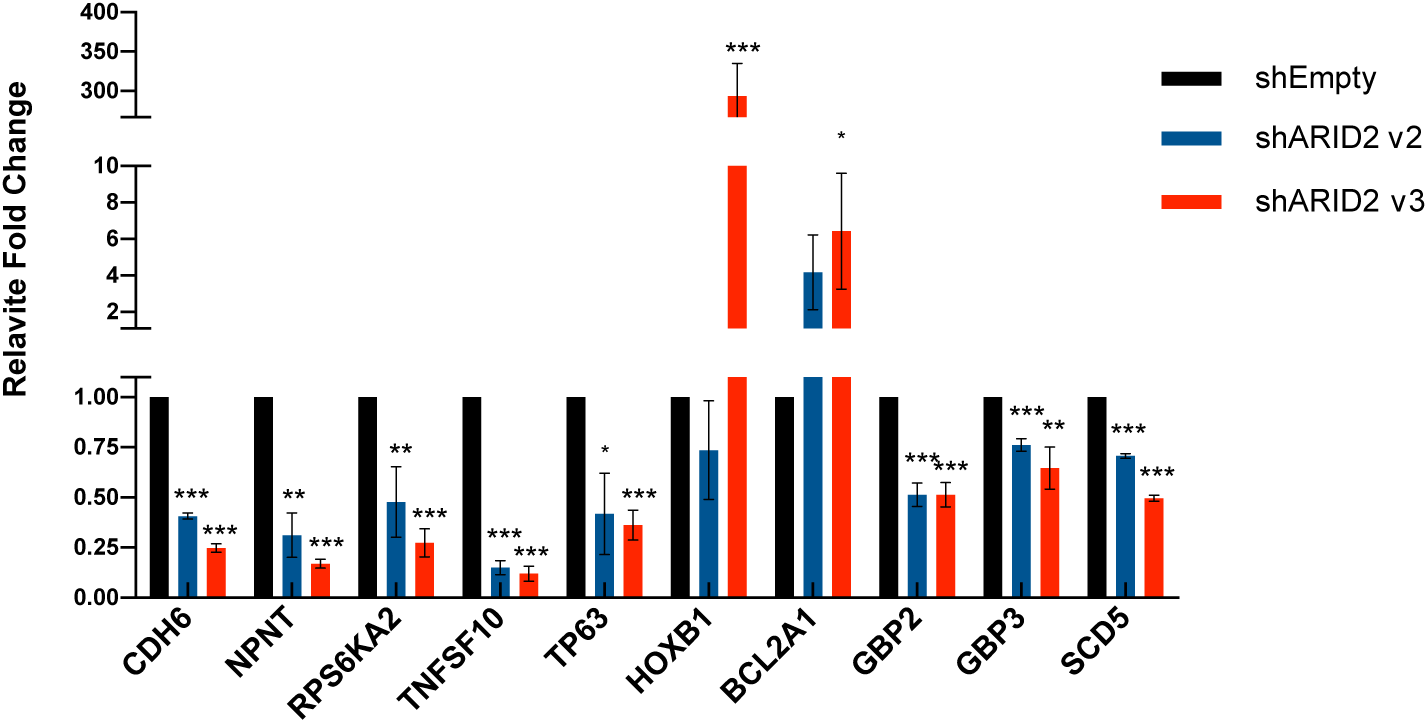
qRT-PCR validation of differential expression changes. Bar graph representation of the results of the qRT-PCR validation of the expression differences identified in selected genes of RNA-Seq experiments. The expression fold-changes are represented as a mean +/- SEM of three independent experiments (one-tailed t-test *p < 0.05, ** p < 0.01 and *** p <0.001).

**Supplementary Figure 5.**
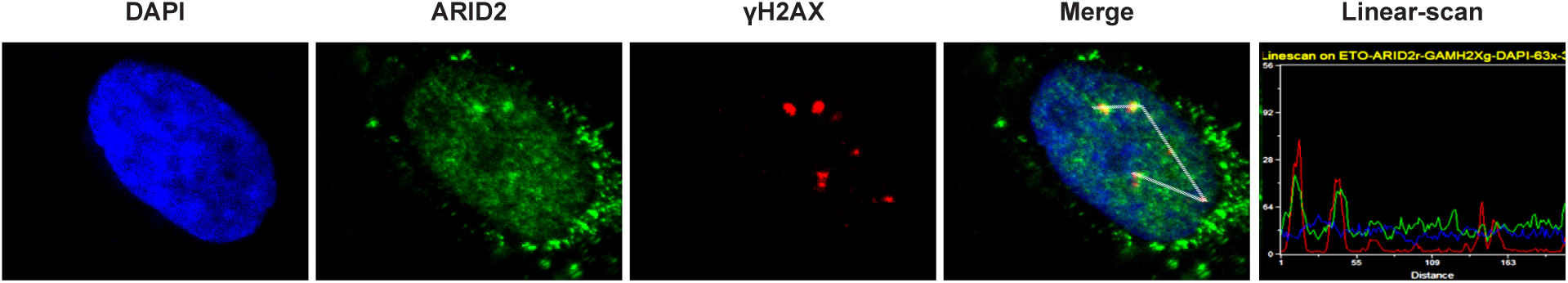
ARID2 co-localizes with DNA repair proteins. Representative images of immunofluorescence experiments proving the colocalization of γH2AX (red) and ARID2 (green) in A549 cells treated with Neocarzinostatin. The colocalization is confirmed through the parallel quantification of read and green signals on a manually selected path through the image using the LineScan software.

**Supplementary Figure 6.**
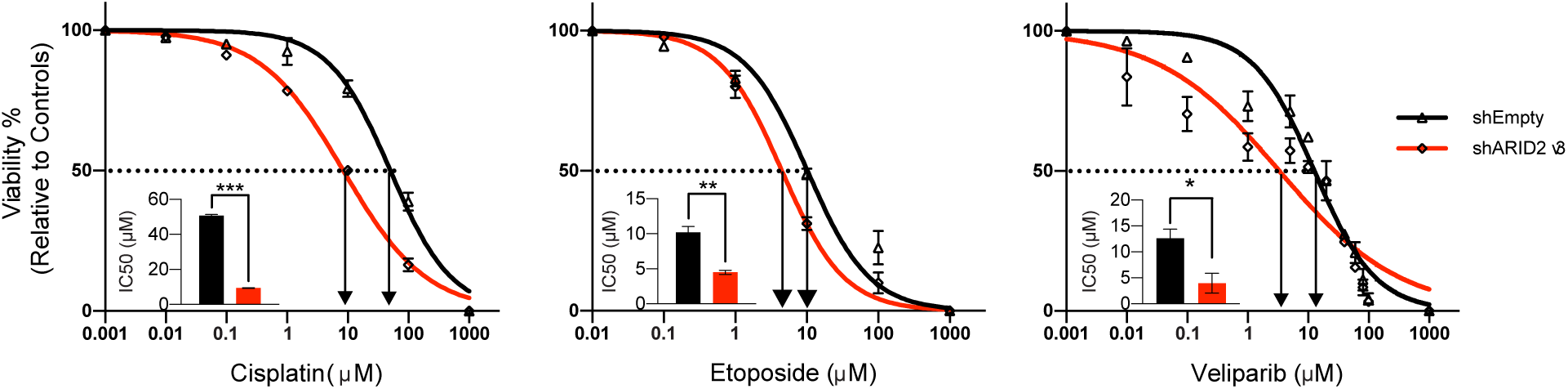
ARID2-deficient cells show different sensitivity to anti-tumor therapies. **R**epresentative experiments measuring cell survival to increasing concentrations of different anti-tumor treatments in H460 cells transduced with shEmpty (black) or shARID2 v3 (red) vectors. Bar graphs represent the calculated IC50 concentrations of H460 cells in each condition. The results are represented as mean +/- SE of three independent results, (one-tailed t-test *p < 0.05, ** p < 0.01 and *** p <0.001).

## Notes

FINANCIAL SUPPORT I.V. is supported by SAF2012-31627 and SAF2016-76758-R grants from the Spanish Ministerio de Economía y Competitividad (MINECO), by a Fundación Ramón Areces grant and by ERC2014-StG637904 grant from the European Research Council. I.V has been awardee of the Programa Ramón y Cajal (MINECO, Spain). T.M has been awardee of the Ayudas para la contratación de investigadores predoctorales (MINECO, Spain). PC laboratory is supported by grant SAF-2015 63638R (MINECO/FEDER, UE); by Centro de Investigación Biomédica en Red de Cáncer (CIBERONC) and by Asociación Española Contra el Cáncer (AECC), grant GCB141423113. BC is supported by a Retos Jóvenes Investigadores grant SAF2015-73364-JIN (AEI/FEDER, UE) and a grant from Fundación Francisco Cobos. P.S. is supported by the Francis Crick Institute, which receives its core funding from Cancer Research UK (FC001152), the UK Medical Research Council (FC001152), and the Wellcome Trust (FC001152).

https://www.ebi.ac.uk/ena/data/search?query=PRJEB26936

