## Supplementary Methods for "*ARID2* deficiency promotes tumor progression and is associated with higher sensitivity to PARP inhibition in lung cancer"

### Patient Samples

Cancer patient primary samples and, when available, matched corresponding normal samples, were obtained from different tumor Biobanks. In all the cases, we counted with the prior approval of the corresponding ethics committee for each institution. A detailed list of the origin and characteristics of each sample can be found in Supplementary Table 1. In total 225 tumor and 87 matched normal DNAs were used in this project. 144 of the tumor samples were sequenced only in the coding region of *ARID2*.

### DNA Extraction and DNA libraries

DNA was extracted from fresh frozen tissue or cell lines using the Agencourt DNAdvance Beckman Coulter kit (#A48705, Beckman Coulter, Brea, CA, USA), following manufacturer's instructions. For the formalin-fixed paraffin-embedded sections, the tumor area was micro-dissected, treated with Proteinase K overnight, subjected to a phenol-chloroform organic extraction followed by ethanol precipitation of the DNA. DNA preparations were quantified using the Qubit® dsDNA BR Assay (Q32851, Life Technologies). Normal DNA libraries were performed mixing from 3 to 5 different DNAs. Diagenode Bioruptor® DNA fragmentation was performed with 500 ng of DNA diluted in low TE buffer (#12090015, Thermo Fisher Scientific, UK) to a final volume of 100 µl, and using 30 cycles of 30''/30'' (ON/OFF cycles) at 4ºC. For all cleaning steps, we used Agencourt AMPure XP (#082A63881, Beckman Coulter, Brea, CA, USA), following the manufacturer's protocol. Size distribution was analyzed with either 2100 Bioanalyzer (DNA 1000 kit) or 4200 TapeStation (D1000 ScreenTape Assay), (Agilent Technologies, Santa Clara, CA, USA). Sequencing libraries were prepared through a series of enzymatic steps including End-repair and Adenylation (DNA Rapid End Repair module, NEXTflex™, #5144-05, Bioo Scientific, Austin, TX, USA), PE adaptor construction through the hybridization of phosphorylated complementary synthetic oligonucleotides, PE adaptor ligation (T4 DNA Ligase, #EL0014 Thermo Fisher

Scientific, UK) and PCR indexing amplification (Phusion high fidelity DNA polymerase, # F530L, Thermo Fisher Scientific, UK). Libraries were checked by Nanodrop for chemical contamination, by the 2100 Bioanalyzer or 4200 TapeStation for size distribution and finally quantified using the Qubit® and a qPCR reaction using primers designed to target the Illumina adapters. Target capture were performed on pools of 96 libraries using a Sure Select® user-defined probe kit (Agilent Technologies, Palo Alto, CA, USA). Massively parallel sequencing was carried out in a High-Seq® machine (Illumina, USA) with a 100bp paired end (PE) protocol. A single lane was performed for each 96-library pool.

In the case of amplicon-based libraries two different strategies were used. Primers were designed for a region of interest using primer3 software (http://primer3.ut.ee/), PCR were performed using 25 to 50 ng as template. Amplified fragments were purified using Agencourt AMPure XP (#082A63881, Beckman Coulter, Brea, CA, USA) following the manufacturer's instructions and submitted to library generation as above. Alternatively, a common Illumina adaptor sequence was included as part of the specific amplification primer. Amplified fragments were purified using Agencourt AMPure XP and submitted to a second PCR for adaptor completion and sample barcoding. Amplicon libraries were sequenced in the MiSeq® platform (Illumina, USA) using a 150 or 250 paired-end protocol according to the amplicon size. A minimum of 50x of average sequence coverage was required for a sample to be processed and included in the study.

**RNA isolation and qRT-PCR.**

Total RNA was isolated and purified using Extract Me Total RNA Kit (Blirt, USA) according to the manufacturer’s instructions. RNA quality was measured using RNA ScreenTape® (4200 TapeStation Instrument - Agilent Genomics). Reverse transcription was performed using the Takara PrimeScript cDNA Synthesis kit (Takara Bio, Inc., Dalian, Japan) according to the manufacturer’s instructions. mRNA expression was measured by qRT-PCR using Luminaris Color HiGreen qPCR Master Mix (Thermo Scientific) with StepOnePlusTM real-time PCR system

(Applied Biosystems, Foster City, CA). β-actin was used as housekeeping gene and the ΔΔCt method was used for quantification and comparison.

**cDNA NGS Libraries**

RNA quality and concentration were measured using a RNA Pico chip on a 2100 Agilent Bioanalyzer. For library preparation, mRNA was enriched using NEBNext® Poly(A) mRNA Magnetic Isolation Module. Fragmentation were performed from 1-2 µg mRNA in a buffer containing 4 µL de PrimeScript Buffer and 1 µL random hexamers primers at 94ºC for 15 minutes. The first strand was synthesized by adding to the previous mix 1 µL of PrimeScript Enzyme and incubating the samples 15 minutes at 37ºC followed by 5 seconds at 85ºC. Second strand was further synthesized by adding to the previous reaction RNAse HI (Thermo) and DNA polymerase I (Thermo) according to manufacturer instructions to a final volume of 100 µL. 2.5 µL de T4 DNA Polymerase (Thermo) and incubated 5 min at 15ºC. 5 µl of EDTA 0.5 M pH 8.0 were added. Fragments were purified using Agencourt AMPure XP (#082A63881, Beckman Coulter, Brea, CA, USA). Library generation protocol was performed starting from the double-stranded cDNA in the same way as DNA libraries, described above. A minimum of 70 million 75 paired-end reads was generated of 6 replicates of each sample group.

**ATAC-Seq Libraries**

ATAC-seq libraries were generated according to Buenrostro et al [1] with minor changes. A549 stably-transduced cell lines were harvested, washed twice in phosphate-buffered saline (PBS), counted. 50,000 cells pellet were resuspended in 50 uL of cold lysis buffer (10 mM Tris-Cl pH 7.4, 10 mM NaCl, 3mM $MgCl_2$, 0.1% IGEPAL, CA-630). Reaction was centrifuged immediately for 10 min at 500 × *g* at 4 °C. Nuclei pellet was resuspended in transposase mixture (25 uL Tagment DNA Buffer, 2.5 uL Tagment DNA Enzyme I (Nextera DNA Library Preparation Kit, Illumina, Cat. 15028220). Nuclei were incubated at 37C for 30 minutes. Transposed DNA was purified using the MinElute Reaction Cleanup Kit (QIAGEN, Hilden, Germany) and eluted in 10 uL of Elution Buffer and and PCR indexing amplification was follow using Phusion high fidelity DNA polymerase kit, (Thermo Fisher Scientific, UK, Cat. F530L) and customized Nextera PCR primers from Buenrostro et al., and the following thermal cycles: 5 minutes at 98 ºC, followed by 15 cycles [98 ºC for 10 s, 62 ºC for 30 s and 72 ºC for 30 s] and a final extension at 72 ºC for 5 minutes. PCR products were purified using Agencourt AMPure XP (Beckman Coulter) and eluted in 20 uL of low TE buffer (#12090015, Thermo Fisher Scientific, UK). Size distribution was analyzed with 4200 TapeStation (D1000 ScreenTape Assay), (Agilent Technologies, Santa Clara, CA, USA), quantified with Qubit® and quality control evaluated by qPCR using Illumina universal sequencing primers.

**DNA-Seq Analysis**

Raw sequence data was subjected to quality control using FastQC v0.11.2 (https://www.bioinformatics.babraham.ac.uk/projects/fastqc/) and mapped to the human genome (hg19) using BWA 0.7.3 [2]. Samtools 0.1.18 [3] was used for format transformation, sorting and indexing of the bam files. Picard 1.61 (http://broadinstitute.github.io/picard/) was used to fix and clean the alignment and to mark PCR duplicates reads. Finally, GATK 2.2.8 [4] was used to perform local realignment around indels. Bedtools 2.17 [5] was used to calculate the enrichment statistics and the target coverage. Paired tumor/normal bam files were used to identify putative somatic single variants (SVs) using an in-house written algorithm called RAMSES [6] selecting mutations with a confidence score of at least 2 and mutational frequency higher than 0.05. PINDEL 0.2.4d [7] was used to detect small insertions and deletions (indels) requiring a minimum of 5 independent reads reporting the indel and with no evidence in the control DNA. Potential germline variants were flagged away using 1000 Genomes mutation database with in-house written software. All *ARID2* mutations were validated by PCR amplification coupled with ultrasequencing at a minimum of 10,000 x coverage. Additionally, a similar orthogonal validation of more than 180 mutations randomly picked showed a near 80% of specificity in the mutation calling. Functional consequence of the mutations was annotated using ensembl database v.73 through the Perl API. OncodriveFML software was run to detect genes with evidence of selective pressure from the analysis [8].

**RNA-Seq Analysis**

Paired-end reads from RNA-Seq were aligned using Tophat [9] to the human genome (hg19). Predicted transcripts from Ensembl database were analyzed and transcripts that would lack a CDS start or stop site were filtered out. Differentially expressed genes (DEG) were identified using HTSeq + DESeq [10,11]. These R packages for transcriptome expression profile analysis were used according to the manufacturer's instructions to test for differential expression of RNA transcript levels requiring a minimum of 3 counts for a gene in more than two independent samples and using a threshold of fold change >2 and a qvalue <0.05. DEG were manually reviewed and the final list of DEG was created.

**ATAC-Seq Analysis**

ATAC-Seq reads were aligned against the human genome (hg19) using BWA 0.7.3 [2]. Accessible regions were identified using MACS 2.1.2 [12]. A combined list of all the regions identified in all the samples was used to perform significant region accessibility in ARID2-deficient cells versus control using DESeq2 [13]. Region annotation was performed using ChIPSeeker software [14]. BEDTools[5] was used to estimate the overlapping of the identified regions with ENCODE publicly available data and the results were plotted using deeptools v3.3.1 [15]. Motif enrichment analysis were performed using HOMER[16]. Finally, alignments were visualized using IGV genome browser [17].

**Cell Culture**

Both A549 and H460 lung cancer cell lines were obtained from The Francis Crick Institute common repository, authenticated by STR profiling, and tested for mycoplasma. The two cell lines were maintained in DMEM (Lonza, Verviers, Belgium) and RPMI 1640 (Lonza, Verviers,

Belgium), respectively, supplemented with 10% FBS (HyClone Victoria, Australia), 1% Gentamycin and 1% Ciprofloxacin at 37°C in a humidified atmosphere containing 5% CO2.

**Generation of stably-transduced cell lines**

For stable cell line generation, tetracycline-inducible pTRIPZ constructs V2THS_74399 (v2), V3THS_347660 (v3) were used for ARID2 knockdown (Dharmacon/GE Healthcare, Lafayette, CO, USA). The empty vector (RHS4750) was used as control. Virus production were performed by transfecting HERK293-T/17clone cells (ATCC ) with the pTRIPZ constructs, psPAX2 and pMD2.G plasmids (Addgene) using Fugene HD (Promega Madison, WI, USA). Infected cells were selected with 1μg/ml puromycin for at least 7 days. Induction of the expression of the shRNAs as well as the turbo-RFP marker was performed with 1μg/ml of Doxycycline for at least 5 days before analyzing the effect of ARID2 knock-down on the cells. Turbo-RFP expression on cells transduced with shRNAs or empty vectors was induced with 1μg/ml Doxycycline for 16h. Cells were isolated by FACS based on TurboRFP expression using the FACS-Aria II cell sorter (Becton Dickinson, BD, Franklin Lakes, USA). For proper cell recovery from the sorting process, the cells were collected in tubes containing DMEM supplemented with 50% FBS to prevent the cells from drying out and dying. Cells were seeded in DMEM complete growth medium.

**Proliferation assays**

Growth curve analyses were performed over a period of fourteen days. Cells were seeded in 100 mm plates at a density of 500,000 cells per plate. Every two days, cells were collected, cell number was determined by counting using a hemocytometer and re-seeded at 500,000 cells per plate. All growth curves were performed in triplicate. As a complementary approximation, PrestoBlue® (Thermo Fisher Scientific, UK) assay was used to determine cell viability with a colorimetric method. Cells were harvested, washed twice in Phosphate-buffered saline (PBS), counted and re-suspended in DMEM complete growth medium at a density of $2 \times 10^4$ cells/ml. Cells were cultured in 96-well plates (BD Falcon, Franklin Lakes, NJ) at a seed density of $2 \times 10^3$

cells/well. 10 µl of PrestoBlue solution was added to the wells and the plates were incubated at 37 °C from 1 to 3 hours. After incubation, absorbance was measured using a Multiskan FC Microplate Photometer (Thermo Fisher Scientific, Waltham, MA) with wavelengths set at 540 and 620 nm.

Cell proliferation was also analyzed using the carboxyfluorescein diacetate succinimidyl ester labeling method with the CellTrace™ CFSE Cell Proliferation Kit (Invitrogen, CA, USA). Transduced cell line was synchronized by gradual serum deprivation following the protocol described by Lauand and collaborators [18]. After 50 h of FBS deprivation, the cells were arrested in G0/G1 phase. Cells were harvested, washed twice in Phosphate-buffered saline (PBS), counted and re-suspended in CellTrace CFSE labelling solution. 1 µl of CellTrace™ stock solution was added to each ml of cell suspension for a final working solution of 5 µM at a density of $10^6$ cells/ml and incubated at 37ºC for 20 minutes protected from light. DMEM culture media containing FBS was added (10% v/v) to remove any free dye remaining in the solution. After 5 minutes, labeled cells were washed with PBS and pelleted by centrifugation. Some of these labeled cells were suspended in fresh pre-warmed complete culture medium and were then seeded into 6-well plates at a density of 5 x $10^5$ cells/well. The remaining labelled cells were suspended in PBS and CFSE fluorescence was measured on a MACSQuant® VYB (Miltenyi Biotec) flow cytometer to identify the parental population and the subsequent division peaks during culture. Cells were harvested at defined times and subjected to division peak resolution by flow cytometry. The cell proliferation index was analyzed using MODFIT software (Verity, USA) . Proliferation index was the sum of the cells in all generations divided by the calculated number of original parent cells.

**Migration assays**

*In vitro* cell migration assays were performed by using 8-µm pore size transwell chambers (Corning™ Transwell™ Multiple Well Plate with Permeable Polycarbonate Membrane Inserts, 3422) in 12-well or 24-well plates. For the migration assays, 200,000 or 50,000 cells were added into the upper chamber. Cells were plated in medium without serum, and medium containing 10% FBS in the lower chamber served as the chemo-attractant. After 24 or 36 h incubation (A549 and

H460, respectively), the cells that did not migrate through the pores were carefully removed using cotton swabs. Filters and lower chamber cells were washed with PBS, harvested by treatment with 0.25% trypsin and counted on a hemocytometer. All experiments were performed in triplicate. Filters were fixed in 4% PFA followed by crystal violet staining for microscope visualization.

**Invasion assays.**

For the invasion assays, 50,000 cells in 50 µl of serum-free DMEM were plated on a growth factor-reduced Matrigel (BD Biosciences) pre-coated 8 µm pore transwell chamber and the lower chamber was filled with 600 µl DMEM with 10% FBS. After 48 or 72 h (A549 and H460, respectively), non-invading cells were removed from the top of the transwells using cotton swabs. Invasive cells were quantified by fixing chambers in 4% paraformaldehyde for 10 min and staining with crystal violet for visualization. Filters and lower chamber cells were washed with PBS, harvested by treatment with 0.25% trypsin and counted on a hemocytometer.

***In vivo* tumorigenesis assays**

Animal studies were conducted in compliance with guidelines for the care and use of laboratory animals and were approved by the Ethics and Animal Care Committee of the Universidad de Cantabria. For proliferation assays, A549 stably-transduced cell lines were harvested, washed twice in phosphate-buffered saline (PBS), counted and re-suspended in PBS at a density of $10^7$ cells/ml. Five million cells were subcutaneously injected into the flanks of the 6-8-week-old female nude mice (Athymic Nude-Foxn1nu, Envigo, UK). The animals were treated with 2 mg/ml of Doxycycline for ~25 days in the drinking water supplemented with 1% sucrose, changed every 2-3 days. After the tumors reached the size of ∼0.5 $cm^3$, mice were euthanizing and tumor tissues were harvested for analyses. For metastasis assays, A549 and H460 stably-transduced cell lines were harvested, washed twice in PBS, counted and re-suspended in PBS + 0.1% BSA at a density 5 x $10^6$ cells/ml. 2.5 million of cells were tail injected into 6-8-week-old female nude mice. The animals were treated for ~60 days with 2 mg/ml of Doxycycline in the drinking water for ~60 days supplemented with 1% sucrose, changed every 2-3 days. After two months, mice were euthanizing and tumor tissues were harvested for analyses.

**Western blot analyses**

Cells were washed twice in PBS and lysed in RIPA buffer (50 mM Tris-HCl, pH 8.0, 150 mM NaCl, 1 % NP-40, 1 mM Sodium Orthovanadate, 1 mM NaF) containing Halt protease inhibitors Cocktail (Thermo Scientific, 87786), for 30 minutes on ice. Lysates were sonicated using the Bioruptor® (Dia-genode) for ten cycles (30 s on, 30 s off) at high-power and cleared by centrifugation at 16,000g for 20 min at 4 °C. Protein concentrations were determined by Qubit® Protein Assay (Q33212, Life Technologies). 50 µg of total protein lysate was separated by SDS-PAGE in 8% polyacrylamide gels and transferred to nitrocellulose membranes. Subsequently, membranes were washed with TBS-T (50 mM TRIS + 150 mM Sodium chloride + 0,1% Tween 20, pH 7,4) and blocked using 5% non-fat milk solution as blocking agent in TBS (50 mM TRIS + 150 mM Sodium chloride) for 1 h at RT. Membranes were then incubated with primary antibodies anti-ARID2 (E-3, sc-166117, Santa Cruz) and anti-Actin (I-19, sc-1616, Santa Cruz), diluted 1:200 and 1: 1,000 in TBS-T/5% (w/v) BSA at 4°C overnight, respectively. After careful washing with TBS-T, the primary antibodies were detected by incubating the membranes with donkey anti-mouse or donkey anti-goat secondary antibodies (LI-COR Biotechnology, Lincoln, USA) conjugated to IRDye 800CW (926-32212) or IRDye 680RD (926-68074) respectively at 1: 15,000 dilutions for 45 minutes at room temperature. Finally, antibody signals were visualized using Odyssey Clx imager (LI-COR Biotechnology, Lincoln, USA).

**Proliferation inhibition *in vitro* assays.**

Inhibition assays were performed to determine the half maximal inhibitory concentration (IC50) values for cisplatin, etoposide and veliparib in both A549 and H460 stably-transduced cell lines. Briefly, cells were seeded in 96-well or 384-well plates at 2000 or 500 cells per well in 100 or 25 µL of complete media, respectively, cultured for 24 hours before drug treatment. Drug concentrations range was prepared in 90 or 25 μl of complete media. Cells were treated for 48 hours. Appropriate media and vehicle controls (DMSO) were also added into the media. Viability was determined by adding 10 μl of PrestoBlue® reagent (Thermo Fisher Scientific, UK) or 25 μl of CellTiter-Glo® Luminescent Cell Viability reagent (Promega, 251 Madison, WI, Cat. G7571). For cell viability assays using PrestoBlue, after incubation, absorbance was measured using a Multiskan FC Microplate Photometer (Thermo Fisher Scientific, Waltham, MA) with wavelengths set at 540 and 620 nm. For cell viability assays using CellTiter-Glo® luminicence was measured GloMax®-Multi+ Detection System Instrument (Promega, 251 Madison, WI, Cat. E9032).

In both cases IC50 value for each drug were determined with Prism software (GraphPad, USA) to fit curves to the dose response data.

**Immunohistochemistry analysis**

For ARID2 detection on paraffin sections, antigen retrieval was performed for 32 minutes at 97 ºC in citrate buffer pH 6. Next, samples incubated with 1:300-1:500 anti-ARID2 antibody (abcam ab113283) and developed with HRP-polymer secondary antibodies (Optiview, Roche).

Immunofluorescence was performed in stable cells induced with 1 μg/mL Doxycycline. Cells reach 50–70% confluence on sterile coverslips were rinsed twice with PBS and fixed with 4% paraformaldehyde in PBS for 15 min at room temperature. Coverslips were rinsed tree times with PBS for 5 minutes. Permeabilization was performed with 0.5% Triton X-100 in PBS for 5 minutes at room temperature. The cells were blocked with 3% BSA in PBT (PBS containing 0.05% Triton X-100) and subjected to immunofluorescence staining with ARID2 antibody(E-3, sc-166117, Santa Cruz Biotechnology Inc., Santa Cruz, CA, USA or A302-230A, Bethyl Laboratories, Montgomery, TX, USA, dilution 1:50), anti-phospho-Histone H2A.X Ser139 (γH2AX, clone JBW301, Merck Millipore, Billerica, MA, USA, dilution 1:400), or anti-53BP1 antibody (H-300, sc-22760, Santa Cruz Biotechnology Inc., Santa Cruz, CA, USA, dilution 1:50) for 30 minutes at room temperature in moist chamber. The cover slips were then washed with PBS three times for 5 minutes. Cells were incubated with Alexa labeled secondary antibodies (1:400) for 30 minutes at room temperature in moist chamber protected from light. Cover slides were mounted in VECTASHIELD Antifade Mounting Medium with DAPI (Vector Labs, Burlingame, CA, USA). The cells were finally examined by fluorescence microscopy (Olympus America Inc, Center Valley, PA or Nikon Eclipse 80i upright fluorescence microscope Tokyo, Japan). Colocalization of ARID2 with 53BP1 or γH2AX was performed measuring the variation in intensity across the lines drawn using the linescan tool from MetaMorph® (Molecular Devices, CA, USA). Automatic foci quantification was performed in randomly acquired images, modified the speckle counting pipeline from CellProfiler open-source software [19].
